# Mitochondria morphology governs ATP production rate

**DOI:** 10.1101/2022.08.16.500715

**Authors:** Guadalupe C. Garcia, Kavya Gupta, Thomas M. Bartol, Terrence J. Sejnowski, Padmini Rangamani

## Abstract

Life is based on energy conversion. In particular, in the nervous system significant amounts of energy are needed to maintain synaptic transmission and homeostasis. To a large extent, neurons depend on oxidative phosphorylation in mitochondria to meet their high energy demand (Pekkurnaz and Wang, 2022). For a comprehensive understanding of the metabolic demands in neuronal signaling, accurate models of ATP production in mitochondria are required. Here, we present a thermodynamically consistent model of ATP production in mitochondria based on previous work (Pietrobon and Caplan, 1985; Magnus and Keizer, 1997; Metelkin et al., 2006; Garcia et al., 2019). The significant improvement of the model is that the reaction rate constants are set such that detailed balance is satisfied. Moreover, using thermodynamic considerations, the dependence of the reaction rate constants on membrane potential, pH, and substrate concentrations are explicitly provided. These constraints assure the model is physically plausible. Furthermore, we explore different parameter regimes to understand in which conditions ATP production or its export are the limiting steps in making ATP available in the cytosol. The outcomes reveal that, under the conditions used in our simulations, ATP production is the limiting step and not its export. Finally, we performed spatial simulations with nine 3D realistic mitochondrial reconstructions and linked the ATP production rate in the cytosol with morphological features of the organelles.

**Summary:** In this work, Garcia et al present a thermodynamically consistent model for ATP production in mitochondria, in which reaction rate constants are set such that detailed balance is satisfied. Simulations revealed that ATP production, but not its export, is the limiting step, and simulations with 3D mitochondrial reconstructions linked the ATP production rate in the cytosol with the morphological features of the organelles.

## 2. Introduction

Energy conversion is at the core of cellular function. Finding efficient strategies to transduce energy has driven the evolution of cells and organisms. In eukaryotic cells, respiration occurs in specialized organelles, known as mitochondria. There, metabolic substrates are converted to adenosine triphosphate (ATP), the energy currency of cell (Alberts et al., 2015). Mitochondrial architecture provides an efficient framework for cellular respiration. It is composed of two distinct membranes: the outer membrane (OM) and the inner membrane (IM). The IM is used to generate an electrochemical proton gradient through a series of electron carriers known as the electron-transport chain (ETC) (Figure 1). The energy stored in this gradient drives the synthesis of ATP from adenosine diphosphate (ADP) and inorganic phosphate (Pi), through a multiprotein complex, ATP synthase (Yoshida et al., 2001). The outer membrane plays a central role in controlling the flow of molecules from and to the interior of mitochondria. Although it forms a barrier to large molecules, it contains a large number of proteins, voltage dependent anion channel (VDAC) (Colombini, 2004), that allow the passage of small metabolites and ions. In contrast, the IM is impermeable to molecules; carrier proteins and ion channels are required for molecules to cross this barrier. One of the most relevant carriers is the adenine nucleotide translocator (ANT), which transports ATP and ADP to and from the matrix, the internal region encapsulated by the IM (Figure 1). Overall, this is a membrane-based process that relies on the lipid bilayer of mitochondria to generate the energy required for ATP synthesis.

**Figure 1.**
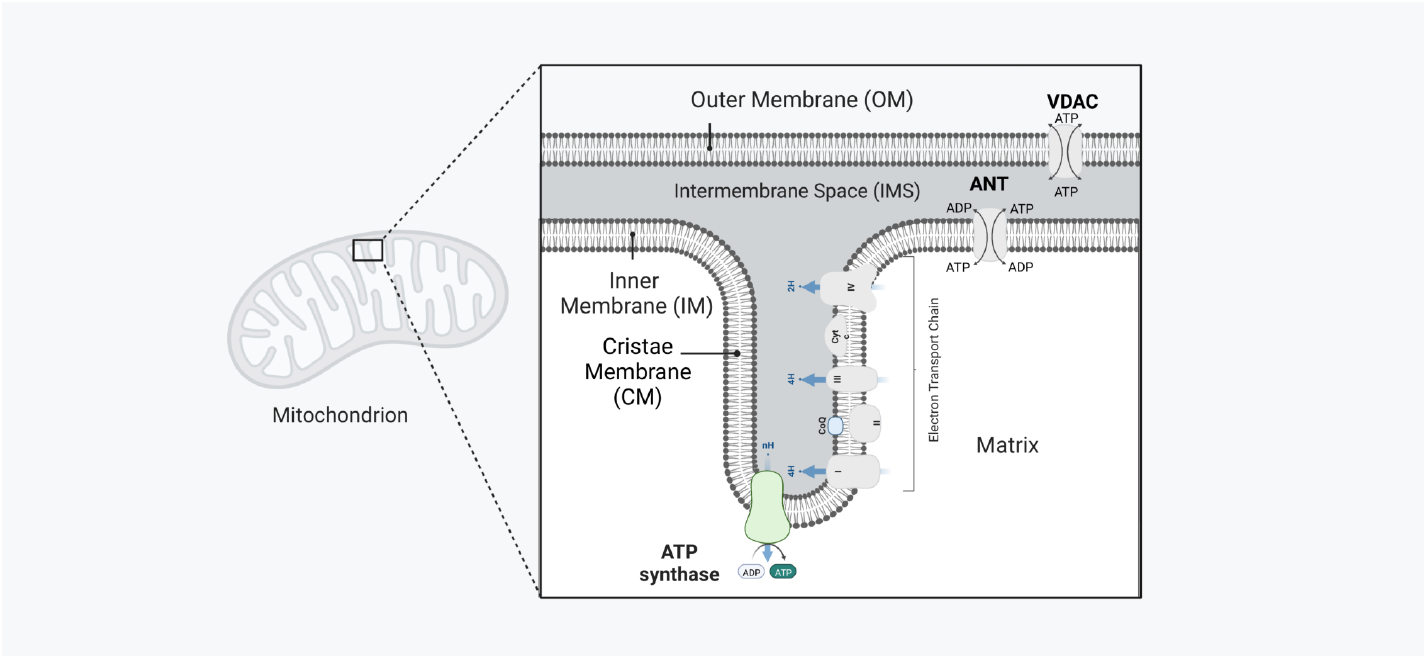
Schematic representation of a mitochondrion with its functional components and compartments. The outer membrane (OM) is the external lipid bilayer that interfaces to the cytosol. The inner membrane (IM) encapsulates the matrix, it is composed of two functionally distinct regions, the inner boundary membrane (IBM) (in close opposition to the OM) and the cristae membrane (CM). The cristae membrane (CM) is the place where the protein complexes from the electron-transport chain (ETC) reside. At the apex of the CM and along the tubular cristae ATP synthases are allocated. Through these protein complexes, ATP is synthesized in the matrix. ATP is further transported from the matrix through adenine nucleotide translocators (ANTs) to the intermembrane space (IMS)—the space between the two membranes— to finally cross the OM at the voltage dependent anion channel (VDAC) into the cytosol.

The importance of ATP production by mitochondria means that reliable mathematical formulations are necessary to build a quantitative framework. To that effect, there have been many mathematical models developed to describe the processes involved in ATP production (Pietrobon and Caplan, 1985; Magnus and Keizer, 1997; Cortassa et al., 2003; Nguyen et al., 2007; Bertram et al., 2006; Beard, 2005; Heiske et al., 2017; Garcia et al., 2019). Pietrobon and Caplan 1985 developed a proton pump model, that can be interpreted as a respiration-driven proton pump or as a model of ATP synthase (Magnus and Keizer, 1997). Later on, Magnus and Keizer 1997 assembled different components and built a steady-state model of ATP production in mitochondria, which several other models are based on (Cortassa et al., 2003; Bertram et al., 2006; Nguyen et al., 2007). However, they did not explicitly show how the thermodynamic constraints were imposed to the system^1^. Some of these models also have been built to be consistent with thermodynamic laws (Beard, 2005; Heiske et al., 2017), and in most cases include more components as the activity of TCA cycle (Cortassa et al., 2003; Nguyen et al., 2007) enzymes or the activity of other components of the ETC (Beard, 2005; Heiske et al., 2017). Almost all previous developed models are implemented with ordinary differential equations (ODEs), disregarding the spatial organization of mitochondria. Here and in our previous work (Garcia et al., 2019) our efforts were focused on developing a minimal reaction-based mitochondrion model so that we can implement it to perform spatial simulations in MCell (Stiles et al., 1996; Kerr et al., 2008; Husar et al., 2022) or non-spatial simulations. In this work, we ensure the model is thermodynamically consistent, for that we imposed detailed balance constraints to the reaction rate constants.

With the model developed here, we answered the following questions: under physiological conditions, what limits the amount of ATP that reaches the cytosol? Is it the rate of ATP production or is it the rate of ATP export from the mitochondria to the cytosol? To answer these questions, we developed a thermodynamically consistent model of ATP production in mitochondria – i.e. a physically plausible model. Our criteria for thermodynamic consistency was to calculate thermodynamic forces (Hill, 1989) for each cycle in the model and to equate them to the ratio of the forward-to-backward reaction rate constants. In this manner, we verified that detailed balance is satisfied for the conditions considered. Furthermore, we calculated the dependence of the reaction rate constants on the membrane potential, pH, ATP, ADP, and Pi concentrations. We conducted simulations of this model using ODEs; although simpler, this approach is informed with detailed structural calculations of surface areas and volumes from a realistic 3D reconstruction (Garcia et al., 2019). The outcomes reveal that, under physiological conditions, ATP production in the matrix is the limiting factor. Lastly, we used the model to perform spatial simulations using nine realistic 3D reconstructions (Mendelsohn et al., 2021). Our results link the ATP production rate to structural features of the mitochondrial geometries.

## 3. Material and Methods

The model presented here is based on previous work (Pietrobon and Caplan, 1985; Magnus and Keizer, 1997; Metelkin et al., 2006; Garcia et al., 2019). Our efforts were focused on assembling a thermodynamically consistent model (Hill, 1989), and explicitly highlighting the dependence of the reaction rates constants on the membrane potential, pH, ATP, ADP, and Pi concentration.

### 3.1. Model for ATP synthase

The ATP synthase model is grounded on the work of Pietrobon and Caplan 1985. The kinetic model consists of a membrane protein –represented as E– that transports protons (H^+^) from the intermembrane space (IMS) to the matrix. The protein can be in six states represented in Figure 2A. Each state has associated a number from 1 to 6 (Figure 2B). A clockwise cycle starting in state 6 (E^−3^) represents the binding of 3 protons from the IMS (transition 6 → 5 in Figure 2B), transport of the protons to the matrix (state 5 → 4), binding of ADP and P_i_ (state 4 → 3) and subsequent synthesis of ATP (state 3 → 2), followed by unbinding of the protons in the matrix (state 2 → 1). We considered a constant proton concentration inside the IMS and the matrix, and the ATP and ADP concentration as variables. The list of the reactions is given in Table 1 and the model parameters are given in Table 2. The number of ATP synthases used in the model was determined employing the density estimated at 3070 µm^−2^ (Acehan et al., 2011). Given the surface area of high curvature of the IM in the reconstructions (Mendelsohn et al., 2021), we estimated the number of ATP synthases at 267 for the mitochondrial reconstruction #1 (Mito 1). We next explain how some kinetic parameters depend on the membrane potential, the proton concentration, and the phosphate concentration.

**Figure 2.**
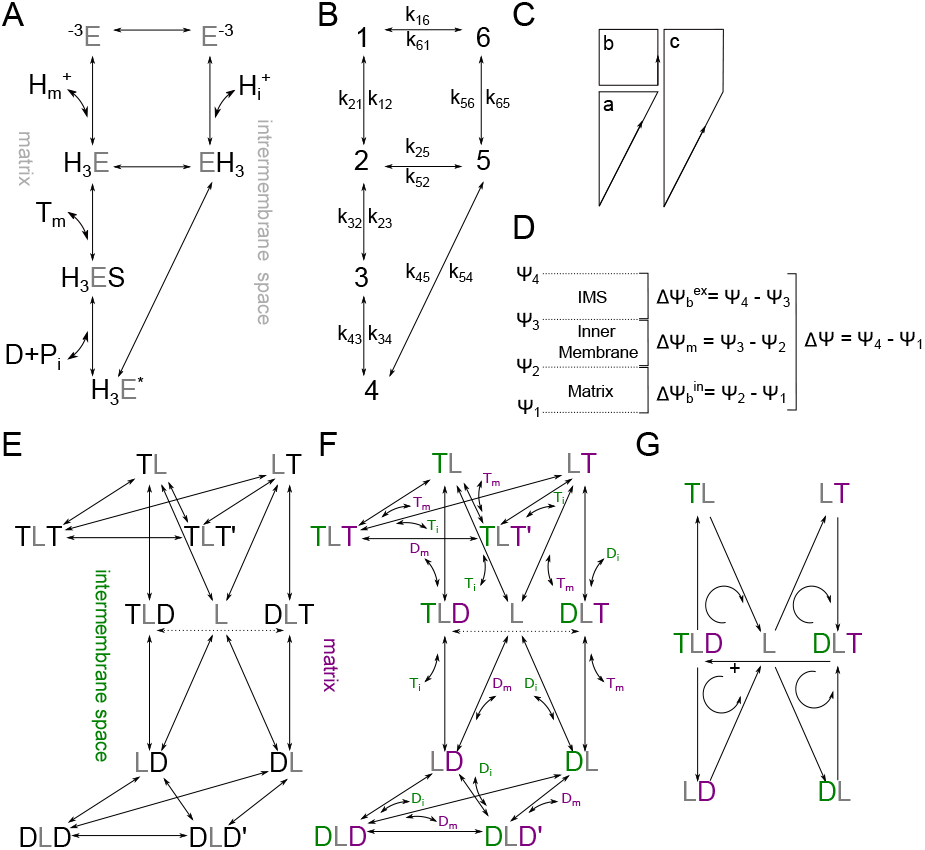
Kinetic diagram of the modules of the mitochondria model. (A-C) Kinetic diagram of the ATP synthase model (Pietrobon and Caplan, 1985). (A) The model consists of six states, representing different protein configurations. ATP molecules are represented as T, ADP molecules as D, Phosphate as P_i_, ATP synthase as E, and H^+^ as protons. The model considers the binding of ATP and ADP from the matrix and the IMS. (B) Each state has associated a number from 1 to 6. A clockwise cycle, starting from state 6 corresponds to the binding of 3 protons from the IMS (6 → 5), followed by their transport to the matrix (5 → 4) and subsequent binding of ADP and P_i_ (4 → 3), to the synthesis of ATP (3 → 2), and unbinding of 3 protons in the matrix (2 → 1). Each transition has an associated rate constant k_ij_. The list of all the reactions is in Table 1 and the list of parameters is in Table 2. (C) The cycles are considered positive in the counter-clockwise direction and named a, b, and c. (D) Membrane potential considered in the ATP synthase model. ΔΨ = Ψ_4_ − Ψ_1_ is the electrical potential difference between the matrix and IMS (or cytosol), ΔΨ_*m*_ is the electrical potential difference across the membrane, 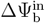 and 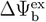 are the phase-boundary potentials. (E)-(G) Kinetic diagram of the ANT model. (E) The model consists of 11 states, L represents the free protein, and YLX is a triple molecular state with one X molecule bound from the matrix side and one Y molecule bound from the IMS. X and Y represent ATP or ADP molecules respectively. The model has been adapted from previous work (Metelkin et al., 2006). (F) In this representation, the binding and unbinding of ATP and ADP are explicitly shown, molecules from the IMS side are in green, and those from the matrix side are in purple. (G) The cycles are considered positive going from the matrix to the IMS.

**Table 1.**
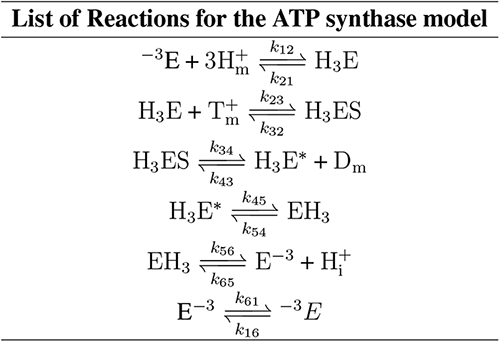
List of reactions for the ATP synthase model.

**Table 2.**
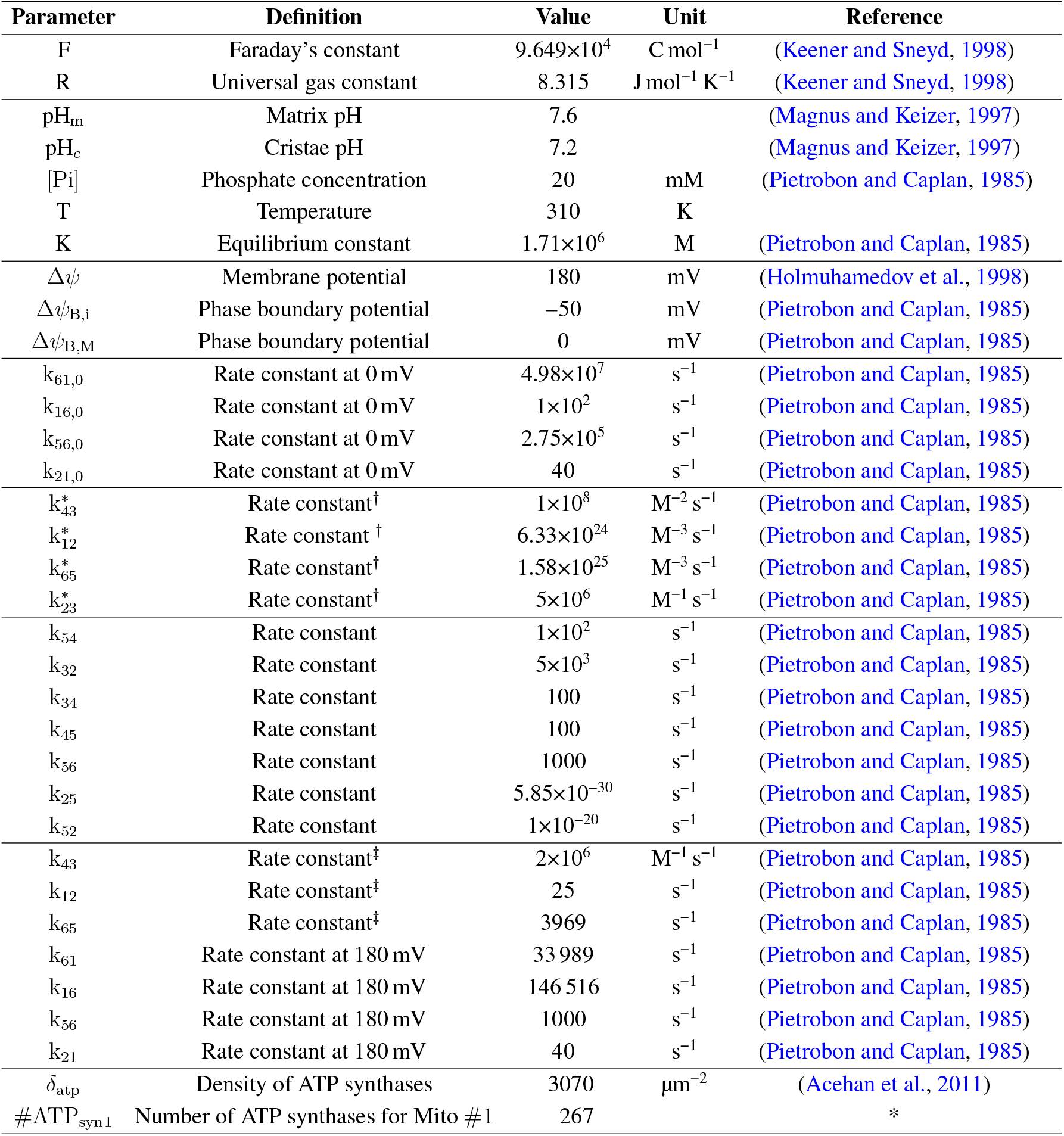
Parameters for the ATP synthase model. * Our estimation, given the surface area of high curvature of the IM measured in the reconstruction (Mendelsohn et al., 2021). † independent of the concentration. ‡ concentration dependent.

#### 3.1.1. Thermodynamic constraints

Thermodynamic constraints are imposed on the rate constants of the system. At an arbitrary steady-state, the product of the first-order (or pseudo-first order) rate constants around a cycle *κ* in the positive direction 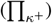 divided by the product of the rate constants in the negative direction 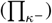 is related to net thermodynamic forces in the following manner (Hill, 1989):

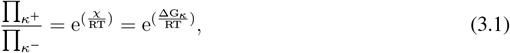

where *χ* is the net thermodynamic force, ΔG_*κ*_ is the difference in Gibbs free energy per mole of substance (i.e. the chemical potential) in the cycle *κ*, R is the universal gas constant and T is the absolute temperature. At thermodynamic equilibrium, and in absence of a potential difference between the membranes, the ΔG in the cycle is zero. This leads to a ratio of forward rate constants to backward rate constants equal to one (Eqs 3.1), also known as detailed balance.

Starting with the shortest cycle, **b** in the diagram (Figure 2C) we obtain:

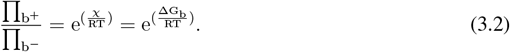

Replacing the rate constants in the cycle we get:

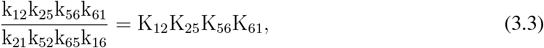

where K_ij_ are the equilibrium constants of the transitions from state i to state j. The equilibrium constants are given by (Caplan and Essig, 1983):

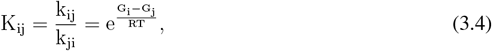

where G_i_ and G_j_ are the Gibbs free energy per mole for species in states i and j respectively. The equilibrium constant for the transition from state 1 to 2, where three protons bind from the matrix side, is equal to:

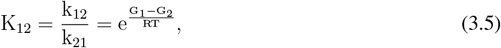

where G_1_ is the Gibbs free energy per mole of three protons in the matrix and G_2_ the energy of the protons bound to the membrane. These are given by (Caplan and Essig, 1983):

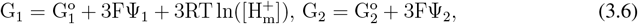

where 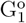 and 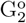 are the standard free energy per mole. Here Ψ_1_ is the bulk potential at the matrix, Ψ_2_ is the phase boundary potential at the surface of the inner membrane (Figure 2D), and F is the Faraday’s constant. Replacing Eq. 3.6 in Eq. 3.5 we arrive at:

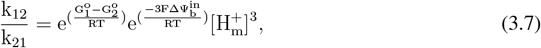

where 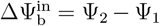 is the phase-boundary potential (the difference between the bulk potential and the potential at the surface of the membrane, see Figure 2D), and 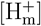 is the proton concentration in the matrix. We define k_ij,0_ as the rate constants at zero potential difference and 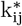 as the rate constants independent of the concentrations; we arrive at:

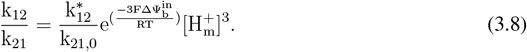

Following the original model (Pietrobon and Caplan, 1985), we assign the effect of the phase-boundary potentials to the unbinding rate constant k_21_, and proton concentration to the binding rate constant k_12_:

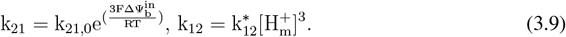

We take a similar approach for the transition 6 → 5, where three protons bind from the IMS to the inner membrane. The Gibbs energy per mole for the protons in the IMS (G_6_) and bound to the membrane (G_5_) are:

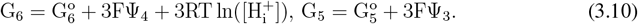

Replacing Eq. 3.10 in Eq. 3.4 we arrive at:

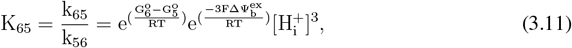

where 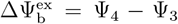 is the phase-boundary potential from the IMS, and 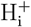 is the proton concentration in the IMS.

As before, we can rewrite Eq. 3.11 with the rate constant at zero potential difference and the rate constant independent of the concentration as:

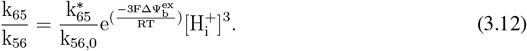

The effect of the phase-boundary potentials is assigned to the unbinding rate constants k_56_, and the proton concentration to the binding rate constant k_65_ and we obtain:

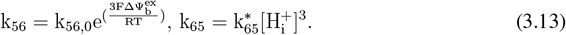

The transition from state 1 → 6 involves the movement of the negatively charged proton biding site from the matrix to the IMS. Therefore the Gibbs free energy for this transition can be written as:

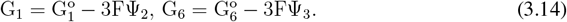

Here, Ψ_2_ and Ψ_3_ are the potentials at the respective membrane surfaces (Figure 2). Replacing in Eq. 3.4 we arrive at:

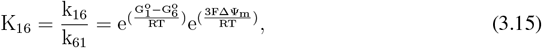

where Δ Ψ_m_ = Ψ_3_ − Ψ_2_. As before, replacing with the rate constants at zero membrane potential difference we reach:

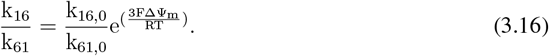

Following the original model (Pietrobon and Caplan, 1985), we considered a symmetrical Eyring barrier (Caplan, 1981). With this assumption the individual rate constants can be written as:

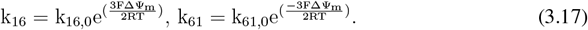

Substituting the equilibrium rate constants calculated above in Eqs. 3.3 we arrive at:

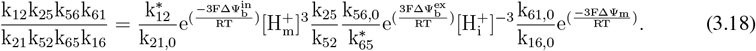

At thermodynamic equilibrium, detailed balance impose an additional constraint to the rate constants:

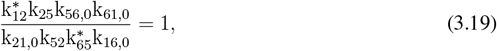

which yields:

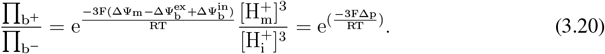

Here Δp is the proton motive force:

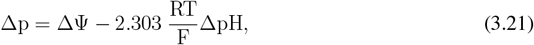

where ΔΨ is the total potential difference across the matrix and the IMS, and ΔpH is the pH difference across the IMS and the matrix (ΔpH = pH_i_ − pH_m_). Similarly for the cycle **a** (Figure 2C) we obtain:

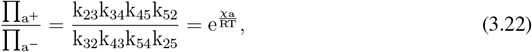

where *χ*_a_ is the thermodynamic force given by:

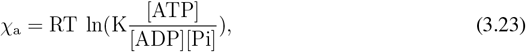

where K is the equilibrium constant of the ATP hydrolysis reaction. At equilibrium, we arrive at:

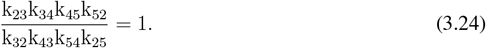

This yields:

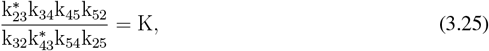

imposing further constraints on the rate constants. Finally, for the longest cycle **c**, the thermodynamic force is the sum of forces for cycles **b** and **a**:

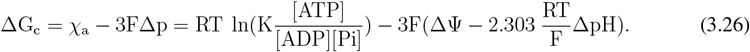

As before the product of the reaction rate constants relates to the thermodynamic forces in the following manner:

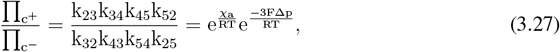

which at equilibrium can be expressed as:

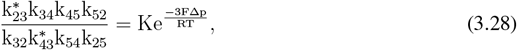

imposing further constraints on the rate constants.

We use the chemical reactions and rate constants presented above to implement the system using compartmental ODEs. As a result of our calculations, the system of ODEs for the ATP synthase model is given by:

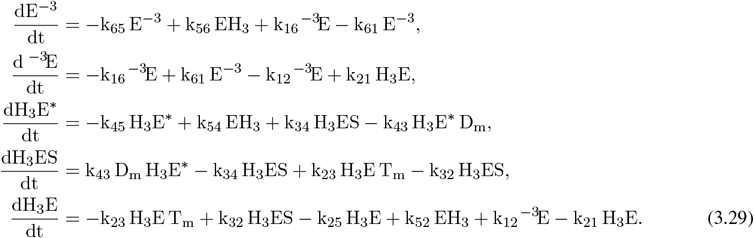

where variables E^−3^,^−3^E, etc, are the number of molecules in the different states. The rate constants of the bimolecular reactions have been normalized to have proper units to compute the number of particles, i.e.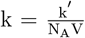, where k^*′*^ represents the constants given in Table 2, N_A_ is the Avogadro’s number, and *V* represents the volume. The volume is either the volume of the matrix or the IMS depending on where the specific reaction occurs (see section 3.5 Simulations Methods). These five differential equations describe the dynamic of the ATP synthase model. The dynamic of the missing state EH_3_ can be deduced, since the total number of proteins is a conserved quantity (E_tot_), leading to the conservation expression E^−3^ + EH_3_ + H_3_E + H_3_ES + H_3_E^*^ + ^−3^E = E_tot_. E_tot_ is the total number of ATP synthases considered in the simulations, this parameter is #ATP_syn1_ in Table 2.

### 3.2. Model for ATP/ADP translocator (ANT)

The ANT model is based on the work of (Metelkin et al., 2006). It is composed of 11 states and 20 bidirectional transitions. The kinetic diagram is shown in Figure 2F. The free protein (L) can bind molecules from the matrix (on the right) or IMS side (on the left), forming a triple molecular state (YLX), where Y and X represent ATP or ADP. State YLX can transition to state XLY exporting X to the IMS and importing Y, following unbinding of the molecules. The productive transition that exports ATP is DLT → TLD, and the reversal reaction imports ATP to the matrix and exports ADP to the IMS. Two additional states (DLD’ and TLT’) were added to track futile translocation (exchange of ADP for ADP and ATP for ATP). Starting from previously fitted flux parameters (Metelkin et al., 2006) for ANTs extracted from heart mitochondria (Kraemer and Klingenberg, 1982), we first estimated the parameters of the dynamic implementation of the model. With this set of parameters, we qualitatively reproduced the independent data from published work (Kraemer and Klingenberg, 1982; Duyckaerts et al., 1980). The rate constants of the model are presented in Tables 4 and 5, and the qualitative reproduction of the experimental data is in Supplementary Figure 1 and Supplementary Figure 2.

**Table 3.**
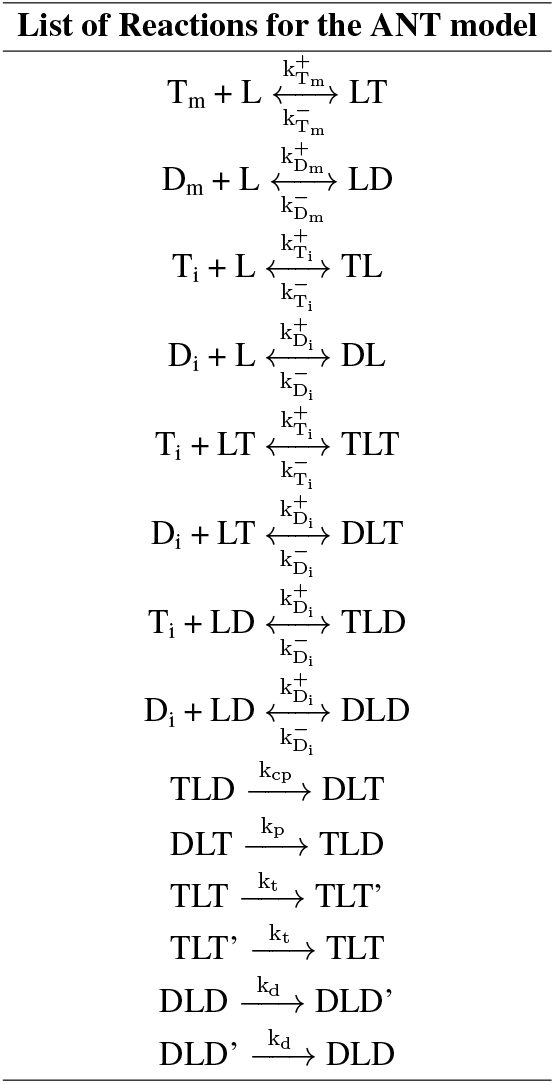
List of Reactions for the ANT model.

**Table 4.** Parameters for the ANT model at Δ*Ψ* = 180 mV. * We estimated 1 nmol/mg ≈ 1.25 mM. ** We used the volume of the IM to get the final number of ANTs.

| Parameter | Value | Unit |
| --- | --- | --- |
| $k_{T_m}^-$ | $4 \times 10^4$ | $s^{-1}$ |
| $k_{T_m}^+$ | $6.4 \times 10^6$ | $M^{-1} s^{-1}$ |
| $k_{T_i}^-$ | 200 | $s^{-1}$ |
| $k_{T_i}^+$ | $4 \times 10^5$ | $M^{-1} s^{-1}$ |
| $k_{D_m}^-$ | $4 \times 10^4$ | $s^{-1}$ |
| $k_{D_m}^+$ | $4 \times 10^6$ | $M^{-1} s^{-1}$ |
| $k_{D_i}^-$ | 100 | $s^{-1}$ |
| $k_{D_i}^+$ | $4 \times 10^6$ | $M^{-1} s^{-1}$ |
| $k_p$ | 9.2 | $s^{-1}$ |
| $k_{cp}$ | 0.35 | $s^{-1}$ |
| $k_d$ | 0.48 | $s^{-1}$ |
| $k_t$ | 0.58 | $s^{-1}$ |
| $T$ | 310 | K |
| $\Delta\psi$ | 180 | mV |
| $\delta_{ANTs}$ | 1.37 | nmol/mg prot* |
| $\#ANTs_1$ | 16471 ** | |

**Table 5.** Parameters for the ANT model at Δ*Ψ* = 0 mV.

| Parameter | Value | Unit |
| --- | --- | --- |
| $k_{T_m}^-$ | $4.4 \times 10^4$ | $s^{-1}$ |
| $k_{T_m}^+$ | $6.4 \times 10^6$ | $M^{-1} s^{-1}$ |
| $k_{T_i}^-$ | 200 | $s^{-1}$ |
| $k_{T_i}^+$ | $4.8 \times 10^6$ | $M^{-1} s^{-1}$ |
| $k_{D_m}^-$ | $4 \times 10^4$ | $s^{-1}$ |
| $k_{D_m}^+$ | $4 \times 10^6$ | $M^{-1} s^{-1}$ |
| $k_{D_i}^-$ | 100 | $s^{-1}$ |
| $k_{D_i}^+$ | $4 \times 10^6$ | $M^{-1} s^{-1}$ |
| $k_p$ | 0.15 | $s^{-1}$ |
| $k_{cp}$ | 0.35 | $s^{-1}$ |
| $k_d$ | 0.48 | $s^{-1}$ |
| $k_t$ | 0.78 | $s^{-1}$ |
| $T$ | 310 | K |
| $\Delta\psi$ | 0 | mV |

#### 3.2.1. Thermodynamic constrains

As presented in the previous section the parameters of the ANT model are constrained by thermodynamic considerations (Eq. 3.1). If we consider a positive cycle a productive translocation of one ATP from the matrix to the IMS and ADP from the IMS to the matrix (Figure 2G), the product of the rate constants in the positive cycle 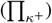 divided by the product of the rate constants in the negative cycle 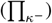 can be written as:

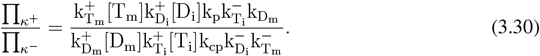

Here k^−^ are the backward rate constants and k^+^ are the forward rate constants for ATP (T) or ADP (D) from the matrix side (m) or the IMS (i), respectively. Replacing with the dissociation constants (the ratio of the backward to forward reaction rates) we arrive at:

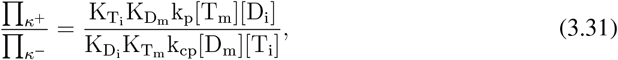

where [D_m_],[T_m_],[T_i_] and [D_i_] are the concentrations of ADP and ATP in the matrix and IMS respectively. The Gibbs free energy per mole associated with the species before the translocation are:

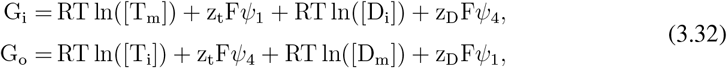

where z_t_ and z_d_ are the charge of the ionized forms of ATP and ADP, −4 and −3 respectively. So the difference in Gibbs free energy per mole associated with the translocation of an ionized ATP molecule from the matrix to the IMS, against a concentration gradient and the translocation of an ionized ADP molecule from the matrix to the IMS is equal to:

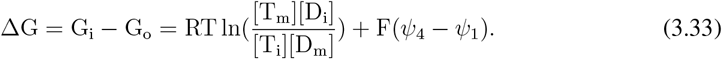

Combining Eq. 3.31 and Eq. 3.33 we arrive at:

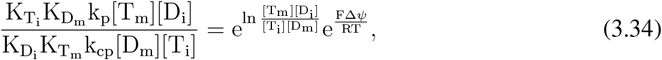

which reduces to:

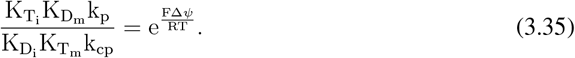

This equation summarizes the imposing restrictions on the rate constants.

The differential equations for the number of molecules of the ANT model in the different states are:

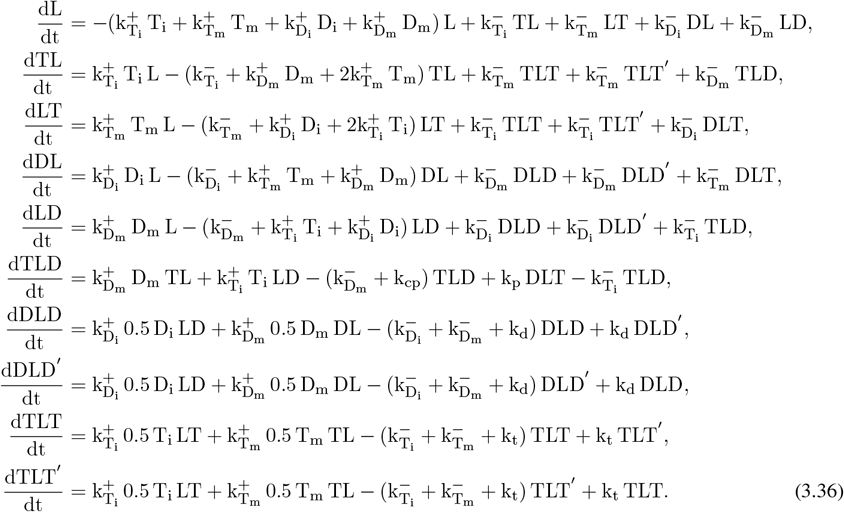

A vast majority of the models do not consider ATP and ADP concentrations as variables (Magnus and Keizer, 1997; Cortassa et al., 2003). Because we focused on the local cytosolic environment around a single mitochondrion, we consider ATP and ADP concentrations in the different compartments as variables. To account for their dynamics, we also derived the equations for the number of ATP and ADP molecules in the different compartments: here D_m_ is the number of ADP molecules in the matrix, T_m_ is the number of ATP molecules in the matrix, D_i_ is the number of ADP molecules in the IMS, T_i_ is the number of ATP molecules in the IMS, T_cyto_ is the number of ATP molecules in the cytosol. These numbers were set by converting the concentration in each compartment to the number of molecules for each reconstruction (see subsection Metabolite buffers 3.4), these were the initial conditions considered. The number of VDACs on the OM is represented as n_vdac_ in these equations (this parameter is #VDACs_1_ in Table 6 for reconstruction #1).

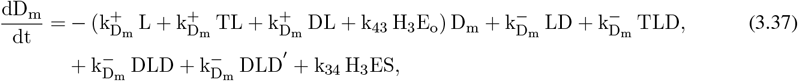

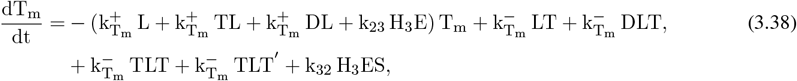

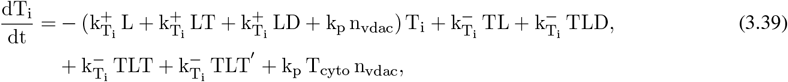

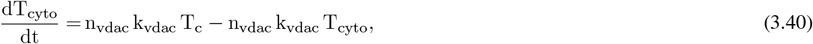

**Table 6.** Parameters for the VDAC model.

| Parameter | Definition | Value | Unit |
| --- | --- | --- | --- |
| $k_{vdac}$ | Rate constant of the reaction | $1 \times 10^6$ | $M^{-1} s^{-1}$ |
| $\delta_{vdacs}$ | Density of VDACS on the OM | $1 \times 10^4$ | $\mu m^{-2}$ (De Pinto et al., 1987) |
| $\#VDACs_1$ | Number of VDACS for Mito # 1 | 6340* | |

As before, the rate constants of the bimolecular reactions have been normalized to have proper units to compute the number of particles, i.e. 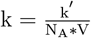, where k^*′*^ represents the constants given in Table 4, N_A_ is the Avogadro’s number, and V represents the volume.

As for the ATP synthase model, the total number of ANTs is a conserved quantity (L_tot_ = L + DL + DLD + DLD^*′*^ + TL + LT + TLT + TLT^*′*^ + DLT + TLD). L_tot_ is the total number of ANTs considered in our simulations, for the mitochondrial reconstruction #1 this quantity is 16471 (this parameter is #ANTs_1_ in Table 4).

### 3.3. Molecular VDAC Model

The main mechanism for metabolites to cross the OM is through VDACs. To consider the ATP export from the IMS into the cytosol, we included VDACs. We implemented a rather simple model assuming VDAC proteins interact with ATP molecules and translocate them to the cytosol by the reaction VDAC + ATP_IMS_ ⇌ VDAC + ATP_cyto_. The list of parameter values employed is given in Table 6.

### 3.4. Metabolite buffers

The total concentration of ATP and ADP can be distributed in different compounds or states like ATP^4−^, ADP^3−^, ATPMg^2−^, etc. ATP and ADP can react with different cations, be bound, or ionized. The final distributions can be estimated by coefficients representing the fraction of unbound ATP in the matrix or the other compartments, these proportions are determined mainly by the level of Mg^2+^. For our model, mitochondrial ADP^3−^ and ATP^4−^ concentrations were estimated analogously to previous work (Magnus and Keizer, 1997). These estimations are based on experimental data (Corkey et al., 1986). The approximate distributions are [ADP]_m,free_ = 0.8 [ADP]_m_, [ATP]_m,free_=[ATP]_m_, [ATP^4−^] = 0.05 [ATP]_free_ and [ADP^−3^] = 0.45 [ADP]_free_.

The initial concentrations of ATP and ADP in the IMS and cytosol were set to 6.5 mM and 0.1 mM (Mörikofer-Zwez and Walter, 1989; Cunningham et al., 1986). The initial concentrations of ATP and ADP in the matrix were set to 13 mM and 2 mM (15 mM in total). The total concentration of adenylates in the matrix (ATP + ADP) of liver mitochondria is in the range 2-27 mM (Rulfs and Aprille, 1982; Nosek et al., 1990; Austin and Aprille, 1984), in the same range for heart mitochondria (Livingston et al., 1996) and also approximately 15 mM for pancreatic cells (Magnus and Keizer, 1997). Moreover, the ratio of ATP to ADP in the matrix has been estimated to be about 10 (Nicholls, 2013). These results are consistent with our assumption of an initial concentration of 13 mM of ATP and 2 mM of ADP in the matrix. In some simulations, these concentrations were kept constant. The concentrations were converted to number of molecules using the volumes of the different compartments (see Table 7).

**Table 7.** Volumes and surface areas of the different compartments considered in the model. These measurements were taken from nine realistic 3D reconstructions of presynaptic and axonal mitochondria (Mendelsohn et al., 2021). The volume of the cytosol (Cyto Vol) is an arbitrary volume. The numbers of proteins considered for each simulation were determined using the densities (Tables 2,4, and 6) and the respective surface areas or volumes. To determined the number of ATP synthases the areas of high curvature (Mendelsohn et al., 2021) were considered (high curva IM SA), values of the first principal curvature larger than 70 *µ*m^−1^ were assessed.

| Property | Mito 1 | Mito 2 | Mito 4 | Mito 5 | Mito 8 | Mito 9 | Mito 10 | Mito 11 | Mito 12 |
| --- | --- | --- | --- | --- | --- | --- | --- | --- | --- |
| OM Vol ($\mu\text{m}^3$) | 0.037 | 0.029 | 0.012 | 0.012 | 0.014 | 0.006 | 0.016 | 0.009 | 0.021 |
| IM Vol ($\mu\text{m}^3$) | 0.016 | 0.010 | 0.005 | 0.004 | 0.006 | 0.002 | 0.008 | 0.003 | 0.010 |
| IM SA ($\mu\text{m}^2$) | 1.545 | 1.104 | 0.505 | 0.390 | 0.508 | 0.224 | 0.605 | 0.314 | 0.806 |
| high curva ($\mu\text{m}^2$) | 0.087 | 0.049 | 0.037 | 0.015 | 0.042 | 0.032 | 0.067 | 0.023 | 0.115 |
| Cyto Vol ($\mu\text{m}^3$) | 0.306 | 0.061 | 0.094 | 0.048 | 0.101 | 0.050 | 0.108 | 0.111 | 0.148 |
| # ATP<sub>syn</sub> | 267 | 150 | 114 | 46 | 129 | 98 | 204 | 72 | 353 |
| # ANT<sub>s</sub> | 16471 | 10294 | 5398 | 4118 | 6177 | 2570 | 7905 | 3578 | 10294 |
| # VDACS | 6340 | 5990 | 2816 | 3370 | 3560 | 1788 | 3126 | 2558 | 3970 |

### 3.5. Simulation methods

#### System of ODEs

The system of ODEs has a total of 19 equations, and 41 parameters, including chemical rate constants, volumes, and number of proteins. The volumes of the different compartments are listed in Table 7, we used the reconstruction #1 for all the non-spatial simulations.

Equations 3.29, 3.37 and 3.40 were integrated using a Radau stiff integrator in PyDSTool (Clewley et al., 2007).

#### Spatial simulations

To perform spatiotemporal simulations in realistic 3D reconstructions we implemented the set of reactions in MCell version 4.0 (Kerr et al., 2008; Stiles et al., 1996; Husar et al., 2022), an agent-based reaction-diffusion simulator. The model is built giving the set of reactions described in Tables 1 and 3. Previous research showed that ATP synthases accumulate at the rim of the cristae membrane in lamellar cristae and along the length of tubular cristae (Strauss et al., 2008; Davies et al., 2012), to emulate this organization in our reconstructions we selected the areas of high curvature in the cristae membrane (Mendelsohn et al., 2021) and we randomly distributed them in these areas. The areas of high curvature were identified as vertices of the cristae membrane with first principal curvature higher than 70 µm^−1^. The location of ANTs in mitochondria has not been definitively resolved, experimental evidence shows they may form complexes with ATP synthases (Ko et al., 2003) located in the CM (Wittig and Schägger, 2009; Vogel et al., 2006). Therefore, we randomly distributed ANTs in the areas of high curvature as well. VDACs were homogeneously distributed on the OM. ATP and ADP molecules were homogeneously distributed in the respective compartments. The diffusion coefficient considered for the free forms of ATP and ADP is 1.5×10^−7^ cm^2^ s^−1^.

#### 3.5.1. Supplementary Material

In Figure 1 and Figure 2 in the supplementary material, we compared experimental results from (Kraemer and Klingenberg, 1982; Duyckaerts et al., 1980) to the results obtained with the ANT translocator model. In Figure 3 we show ATP fluxes through ANTs generated with the model developed by (Bohnensack, 1982). In Figure 4 we present spatial simulation performed with the same conditions considered in the main text, but keeping the number of ATP synthases constant in all reconstructions. In Figure 5, we compare the result with the previously developed version of the mitochondrial model (Garcia et al., 2019) with the current version.

**Figure 3.**
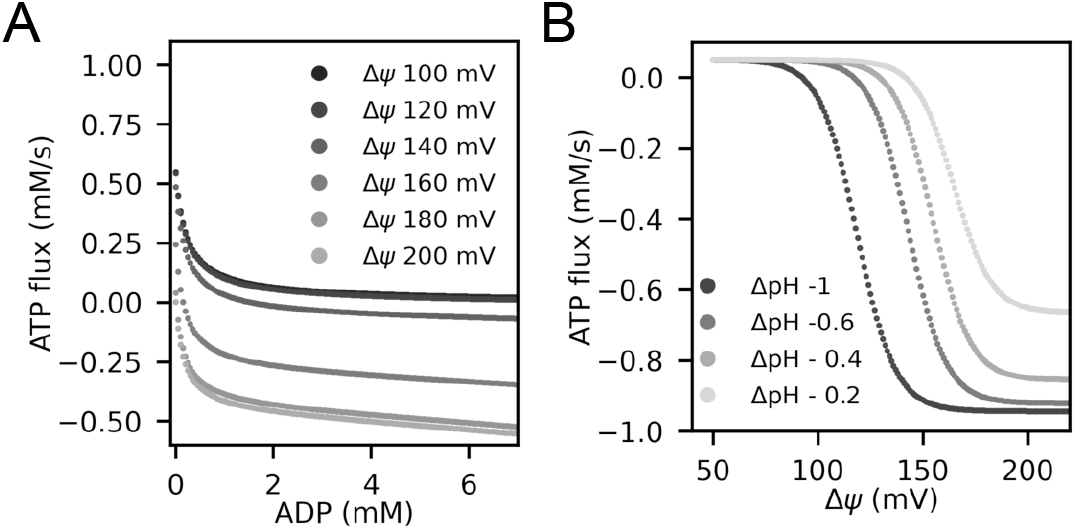
ATP synthase steady-state flux analysis. We analyzed the dependence on the fluxes through ATP synthases at constant concentrations of adenine nucleotides in the different compartments. We analyzed how the flux changes with the driving forces of the cycles (Eqs. 3.26). A) ATP net flux through ATP synthases for different concentrations of ADP in the matrix and different membrane potentials. A counter-clockwise cycle is defined as positive, favoring ATP hydrolysis. For membrane potential above 160 mV and larger ADP_m_ concentrations, ATP is synthesized. B) ATP net flux through ATP synthases for different pH and membrane potentials (for fixed concentrations of adenine nucleotides).

**Figure 4.**
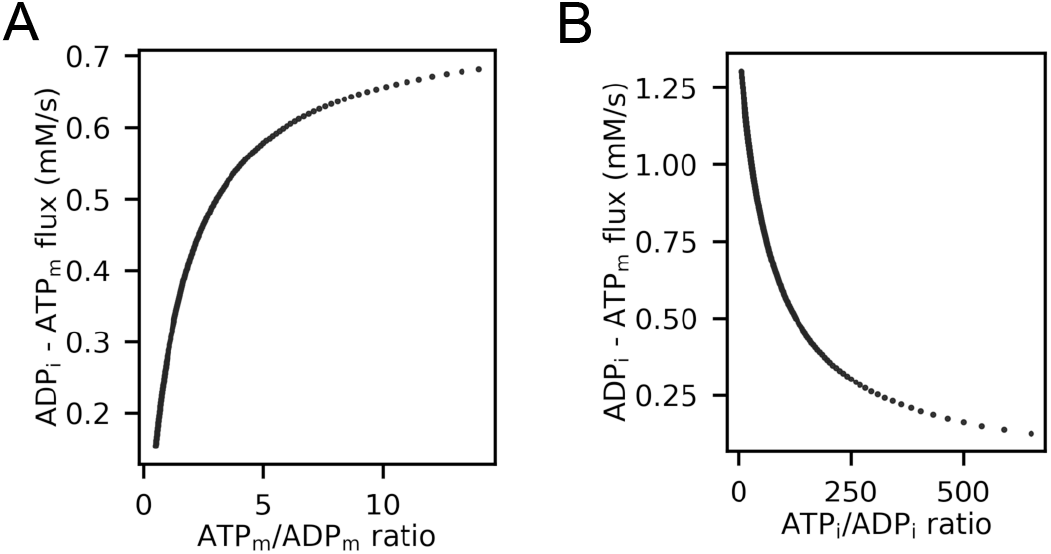
ANT steady-state flux analysis. We analyzed the dependence on the fluxes through ANTs at constant concentrations of adenine nucleotides in the different compartments and at a constant membrane potential of 180 mV. We analyzed how the flux changes with the driving forces of the cycles (Eqs. 3.33). A) ADP_i_ − ATP_m_ net flux through ANTs for different ATP to ADP ratios in the matrix. B) ADP_i_ − ATP_m_ net flux through ANTs for different ratios of ATP and ADP in the IMS.

**Figure 5.**
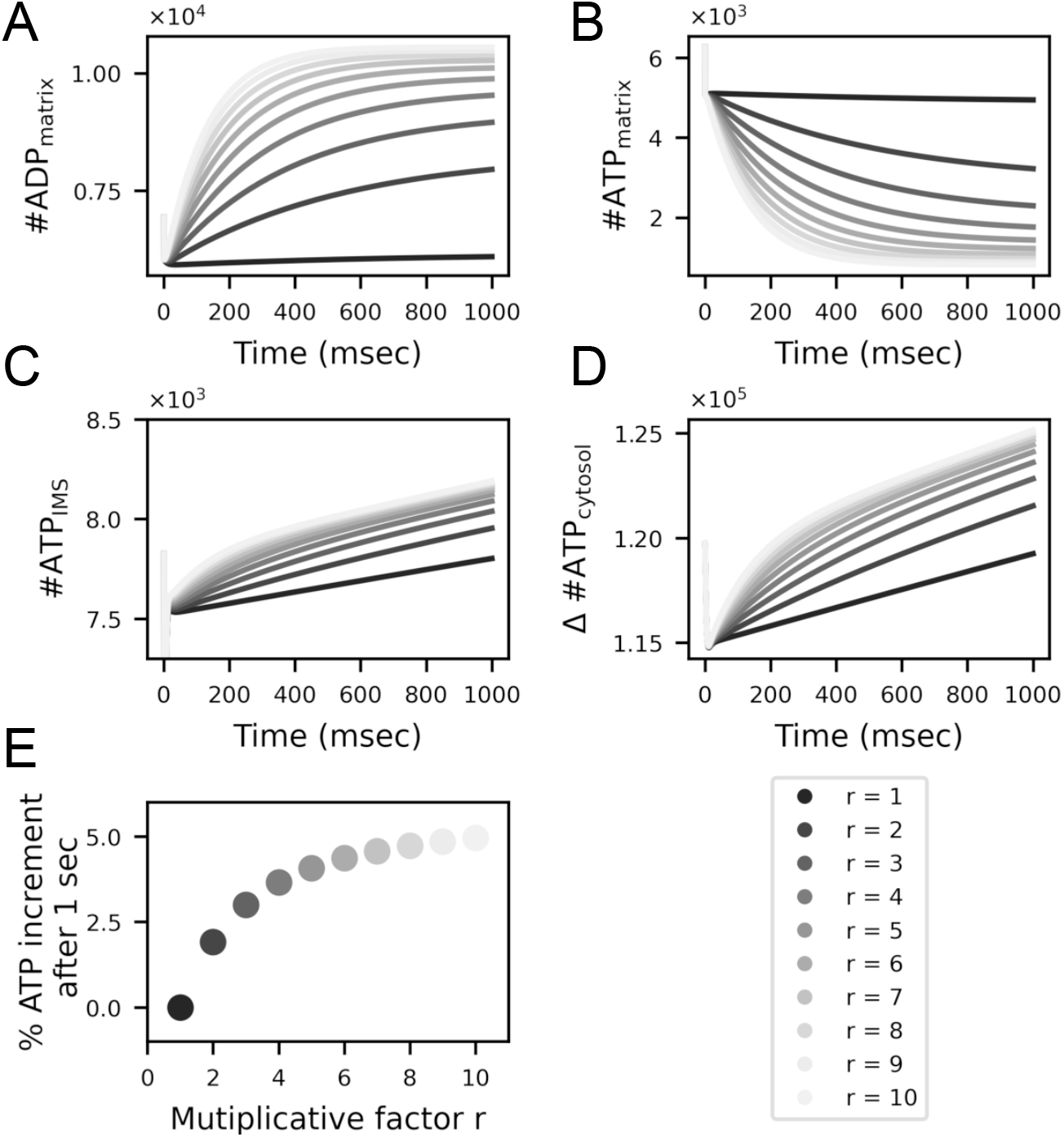
Parameter exploration for the ANT model at a constant membrane potential of 180 mV. We employed the ODE model to explore the impact of faster ANTs on ATP production, for that, we changed the rate constants of the ANTs. We multiplied the rate constants k_p_, k_cp_, k_t_, and k_d_ by a factor r, where r takes values between 1 and 10. The ADP concentration in the IMS was kept constant in these simulations. (A)-(C) Number of molecules measured over time in the different compartments. (D) Number of ATP molecules measured over time in the cytosol relative to the initial number of molecules. (E) Percentage increment of the number of ATP molecules in the cytosol after 1 second relative to the slowest ones (r = 1), for all ANTs considered from the slowest (r = 1) ones to the fastest ones (r = 10).

## 4. Results

Before studying the dynamics of the assembled model, we first analyzed the behavior of its components separately and compared them with previously developed models. In these first simulations, we assumed steady-state conditions and kept the ADP and ATP concentrations constant in the different compartments. We then conducted dynamic simulations with the ODE representation to understand which is the limiting step in ATP production. Finally, we performed spatial simulations with nine realistic 3D geometries and related the ATP production rate to the morphology of the organelles.

### 4.1. Steady-state flux analysis for the model components and comparison to previously developed models

First, we analyzed the dependence of the ATP flux on the ADP concentration, membrane potential, and pH for the ATP synthase model. The driving forces of the cycle **c** (Figure 2C) are the membrane potential, the pH, and the ratio of the ADP and Pi concentration, to the ATP concentration relative to the equilibrium constant (Eq 3.26). For these simulations, the ΔpH was set to -0.4 (Tischler et al., 1977) and the total concentration of adenine nucleotides in the matrix was set to 15 mM (Austin and Aprille, 1984; Rulfs and Aprille, 1982). In Figure 3A, we show the ATP net flux through ATP synthases for different concentrations of ADP in the matrix. A positive flux is defined in the counter-clockwise direction, favoring ATP hydrolysis. We found that at small concentrations of ADP the flux is positive, meaning ATP is hydrolyzed. Increasing the concentration of ADP in the matrix, the ATP flux becomes negative, indicating that ATP synthesis is favored. The flux saturates for concentrations of ADP above 1 mM. Moreover, the driving forces of the cycle also depend on the membrane potential (Eq. 3.26). For a membrane potential above 160 mV the ATP flux is negative, for large concentrations of ADP, but for membrane potential below those values ATP is hydrolyzed. Similar results were obtained for the ATP flux derived by Magnus and Keizer 1997, and later modified by Cortassa et al. 2003 (for comparison see Figure A6-A in the Supplementary Material by Cortassa et al. 2003, the difference in the sign of the fluxes is due to different sign conventions).

Next, we studied the dependence of the ATP flux on the pH difference between the matrix and the IMS (Figure 3B). Here, we fixed the pH in the matrix at 7.6 and varied the pH in the IMS. When the pH difference between the matrix and IMS is large, a smaller membrane potential is needed to observe a negative ATP flux, i.e. to synthesize ATP. This is because the proton motive force —the membrane potential minus the difference in pH (Eq. 3.21)— drives the protons to the matrix.

The ATP flux also saturates at higher membrane potentials, and in particular, it is almost saturated at 180 mV —the potential considered for the rest of this work. Similar flux dependencies were observed in the past (for comparison see Figure A6-B in the Supplementary Material by Cortassa et al. 2003). Although their maximal flux is about 10 mM s^−1^ while ours is about 0.9 mM s^−1^. A similar dependence on the membrane potential was also observed by Nguyen et al. 2007 (Figure 2D in their paper), their maximum flux is 1.125 mM s^−1^ very close to the range in our work.

The driving force of the ANT translocation is the membrane potential and the ratio of ATP_m_*/*ADP_m_ in the matrix and ATP_i_*/*ADP_i_ in the IMS (Eq. 3.33). Therefore, we studied the dependence of the net ADP_i_ − ATP_m_ flux through ANTs at steady concentrations of ATP and ADP, for different ratios. First, we calculated the net ADP_i_ − ATP_m_ flux through ANTs for different ratios ATP_m_*/*ADP_m_ in the matrix (Figure 4A). Keeping the cytosolic ratio fixed at 65, we varied the ATP_m_*/*ADP_m_ ratio. The driving force of the cycle (Eq. 3.33) increases for larger ATP_m_*/*ADP_m_ ratios, increasing the net ADP_i_ − ATP_m_ flux (Figure 4A). The net ADP_i_ − ATP_m_ flux saturates for ratios ATP_m_*/*ADP_m_ closer to the ratio of ATP_i_*/*ADP_i_ (Eq. 3.33).

Next, we varied the ATP_i_*/*ADP_i_ ratio in the IMS and kept the matrix ratio constant at 6.5. Increasing the ATP_i_*/*ADP_i_ ratio decreases the driving force (Eqs. 3.33) at constant ATP_m_*/*ADP_m_ ratio. Therefore, the net ADP_i_ − ATP_m_ flux decreases for larger ratios of ATP_i_*/*ADP_i_ (Figure 4B). Moreover, the maximum flux is set by the ATP_m_*/*ADP_m_ ratio. Similar flux dependencies were found by Magnus and Keizer 1997 with the model developed by Bohnensack 1982, see the Supplementary Figure 3.

From this analysis, we identified that for 1 mM ADP_m_, 180 mV membrane potential, and -0.4 ΔpH, the ATP flux is almost at its maximum absolute value (Figure 3A-B). We used these values for the rest of this work. Likewise, the initial concentration ratios considered in the rest of this work are ATP_m_*/*ADP_m_ = 6.5 and ATP_i_*/*ADP_i_ = 65, from which the ADP_i_ − ATP_m_ flux through ANTs is almost at its maximal value.

### 4.2. Under physiological conditions, ATP production is the limiting step for cytosolic ATP availability

The amount of ATP in the cytosol depends on the production rate and its export through ANTs and VDACs. Since experimental results suggest that the ANT translocation rate constants are in the order of 100s per second (Gropp et al., 1999), we explored the impact of fast export through ANTs on the final ATP concentration in the cytosol. We held the concentration of ADP in the IMS constant and changed the parameters of the ANT model. Specifically, we changed the value of the rate constants k_p_, k_cp_, k_t_, and k_d_ multiplying them by a factor r, such that detailed balance is still satisfied. We measured the number of ADP and ATP molecules in the matrix (Figure 5A-B), and the number of molecules of ATP in the IMS and cytosol (Figure 5C-D). Faster translocation allows more ADP to accumulate in the matrix (Figure 5A) and therefore less ATP (Figure 5B), while ATP molecules reach the IMS and the cytosol faster (Figure 5C-D). We further calculated the percentage of the ATP increase relative to the slowest import (Figure 5E). The fastest ANTs (r = 10) exports 5% more ATP molecules than the slowest ones. Moreover, the amount of ATP starts to saturate for the fastest ANTs (r = 10). This means even ANTs that are faster cannot overcome the ATP production rate. Thus, the rate-limiting step is the ATP production rate, not the export.

In addition to the translocation rate of the ANTs, the ratio of ANTs to the ATP synthases can influence the export of ATP molecules into the cytosol. Therefore, we next asked how this ratio affects the availability of ATP in the cytosol for the fastest ANTs. We asked which step limits the number of ATP molecules in the cytosol under these conditions – is it the synthesis of ATP or its translocation through ANTs? As in the previous simulations, the number of ADP molecules in the IMS was constant, assuming unlimited resources. We measured the number of ADP and ATP molecules in the matrix, IMS, and cytosol over time (Figure 6A-D).

**Figure 6.**
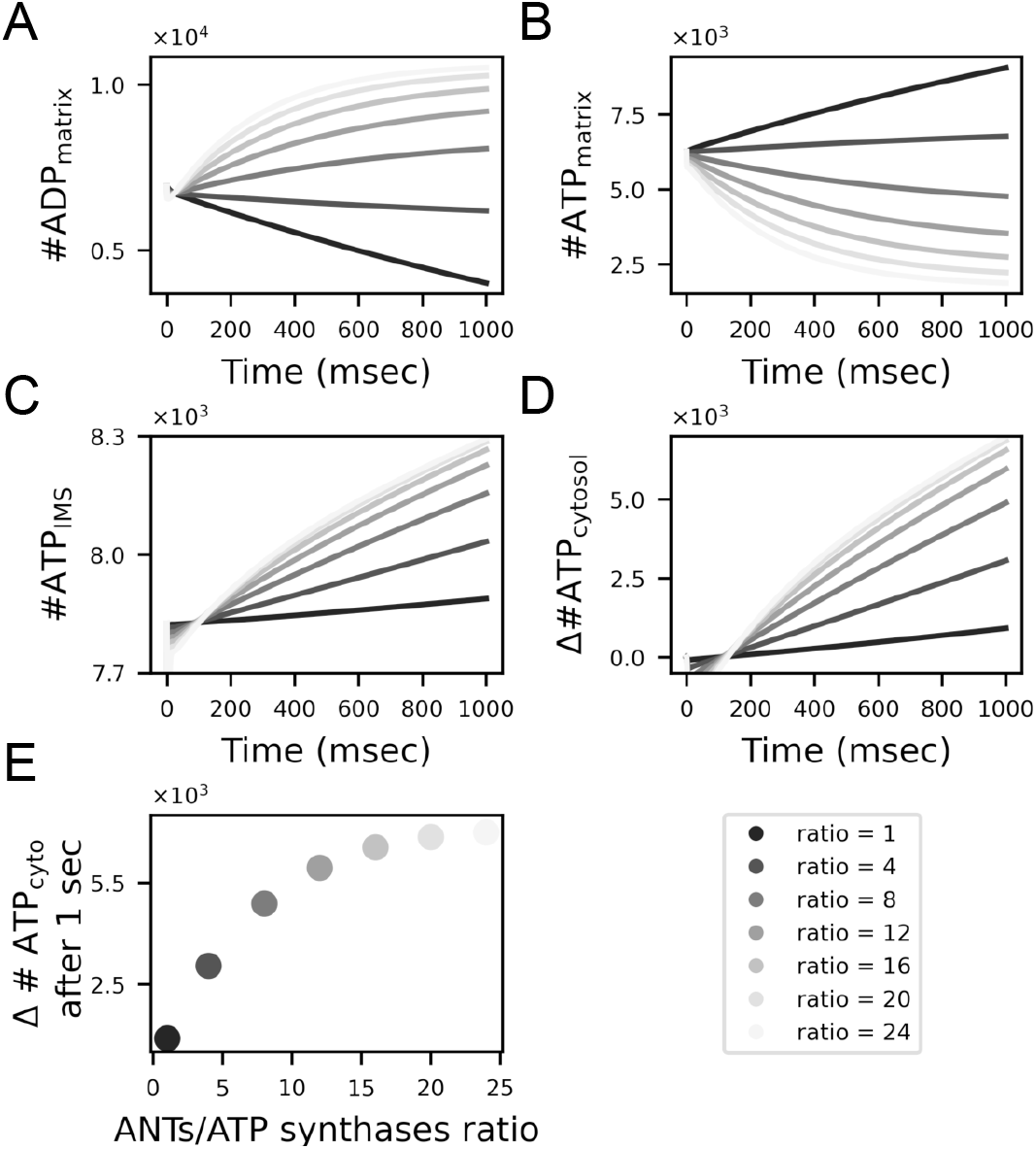
What is the limiting factor, export through ANTs or production of ATP through ATP synthases? We changed the ratio of ANTs to ATP synthases to answer this question and used the ODE model to estimate the impact on ATP production, for a constant concentration of ADP in the IMS and a constant membrane potential of 180 mV. (A)-(C) Number of molecules measured over time in the different compartments. (D) Number of ATP molecules measured over time in the cytosol relative to the initial number. (E) Number of ATP molecules in the cytosol after 1 second relative to the initial number of molecules, for different ratios of ANTs/ATP synthases. Ratios go from 1 to 24, the number of ATP synthases is kept constant and the number of ANTs increases (fast ANTs were considered r = 10).

The amount of ATP in the cytosol after 1 second starts to saturate when the number of ANTs is twenty times the number of ATP synthases (Figure 6E). Thus our model predicts that having more than twenty times ANTs to ATP synthases number will not have any benefit. If this ratio is bigger than twenty, the limiting step is the production of ATP.

Similarly, VDACs can limit the amount of ATP in the cytosol. Therefore, we analyzed how the export of ATP through VDACs limits their amount in the cytosol. Considering similar conditions as before constant concentration of ADP in the IMS, we changed the number of VDACs in the OM and measured the number of ADP and ATP in the different compartments (Figure 7A-D), along with the percentage increment of ATP in the cytosol after 1 second (Figure 7F). Having less than approximately 500 VDACs in the OM significantly limits the amount of ATP available in the cytosol. Otherwise, the production of ATP is the limiting step.

**Figure 7.**
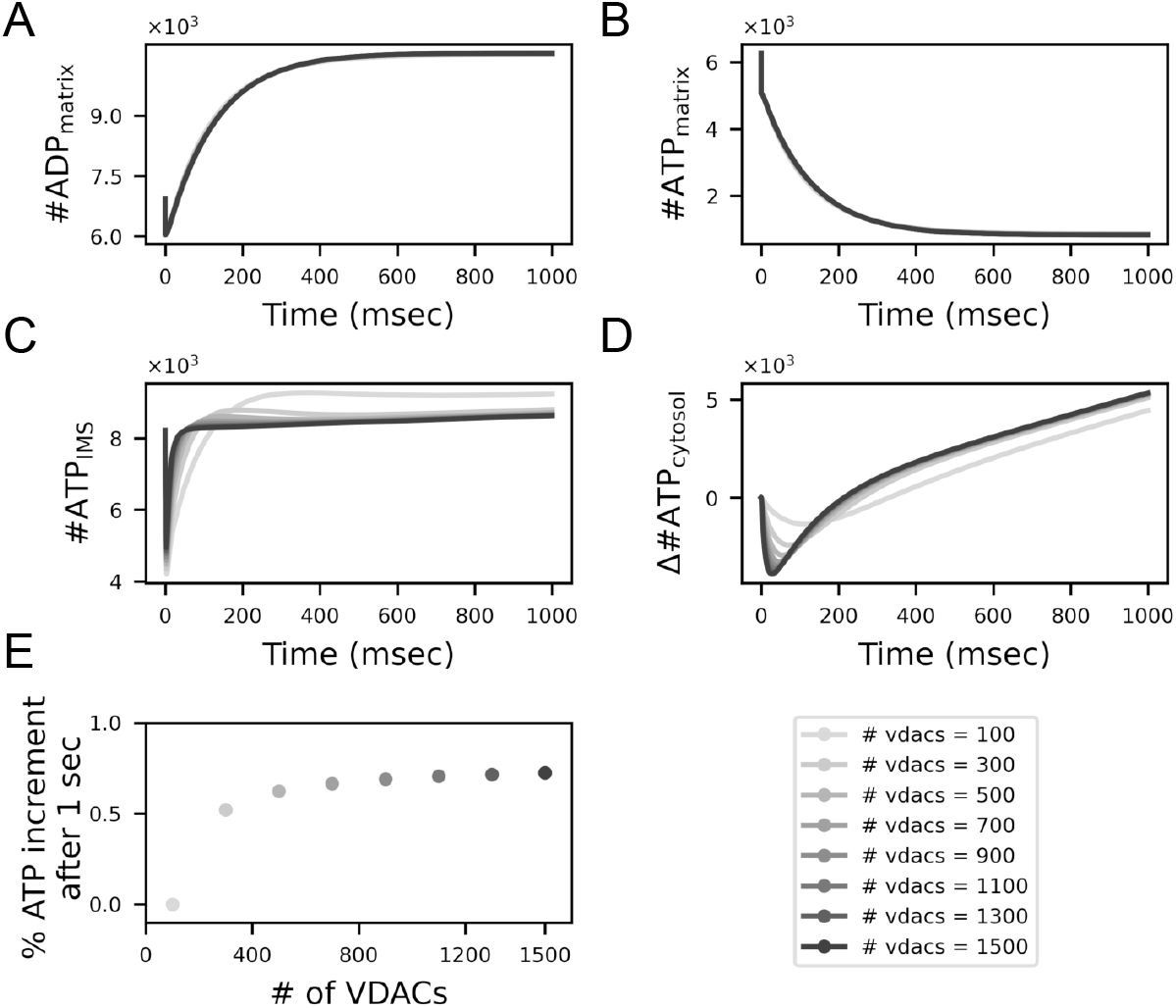
Effect of the number of VDACs on ATP efflux to the cytosol at a constant membrane potential of 180 mV. (A)-(C) Number of molecules measured over time in the different compartments. (D) Number of ATP molecules measured over time in the cytosol relative to the initial concentration. (E) Percentage increase of ATP in the cytosol after 1 second for different numbers of VDACs in the OM. The number of VDACs goes from 100 to 1500 (in these simulations fast ANTs were considered, r = 10).

Given the density of VDACs in the OM (De Pinto et al., 1987) and the surface area of the OM measured (Mendelsohn et al., 2021) (see Methods), the number of VDACs in the reconstruction #1 is 6340. This number is well above the limiting threshold of 500. Therefore, in the physiological conditions we considered, the production of ATP is the limiting step.

### 4.3. Spatial Simulations: application of the model to different 3D mitochondrial reconstructions

To illustrate the utility of our model, we conducted spatial simulations with nine realistic 3D reconstructions (Mendelsohn et al., 2021), using the agent-based reaction-diffusion simulator MCell, and we related the ATP production rate to morphological features of mitochondria. In Figure 8A we show the CM of the nine reconstructions with the total number of ATP synthases considered for the simulations (in red). In Figure 8B we show the results of the ODE system and averaged traces of the spatial simulations (over ten different realizations) of the number of ADP and ATP in the different compartments for the elongated mitochondrial reconstruction #2, for the same conditions considered as before.

**Figure 8.**
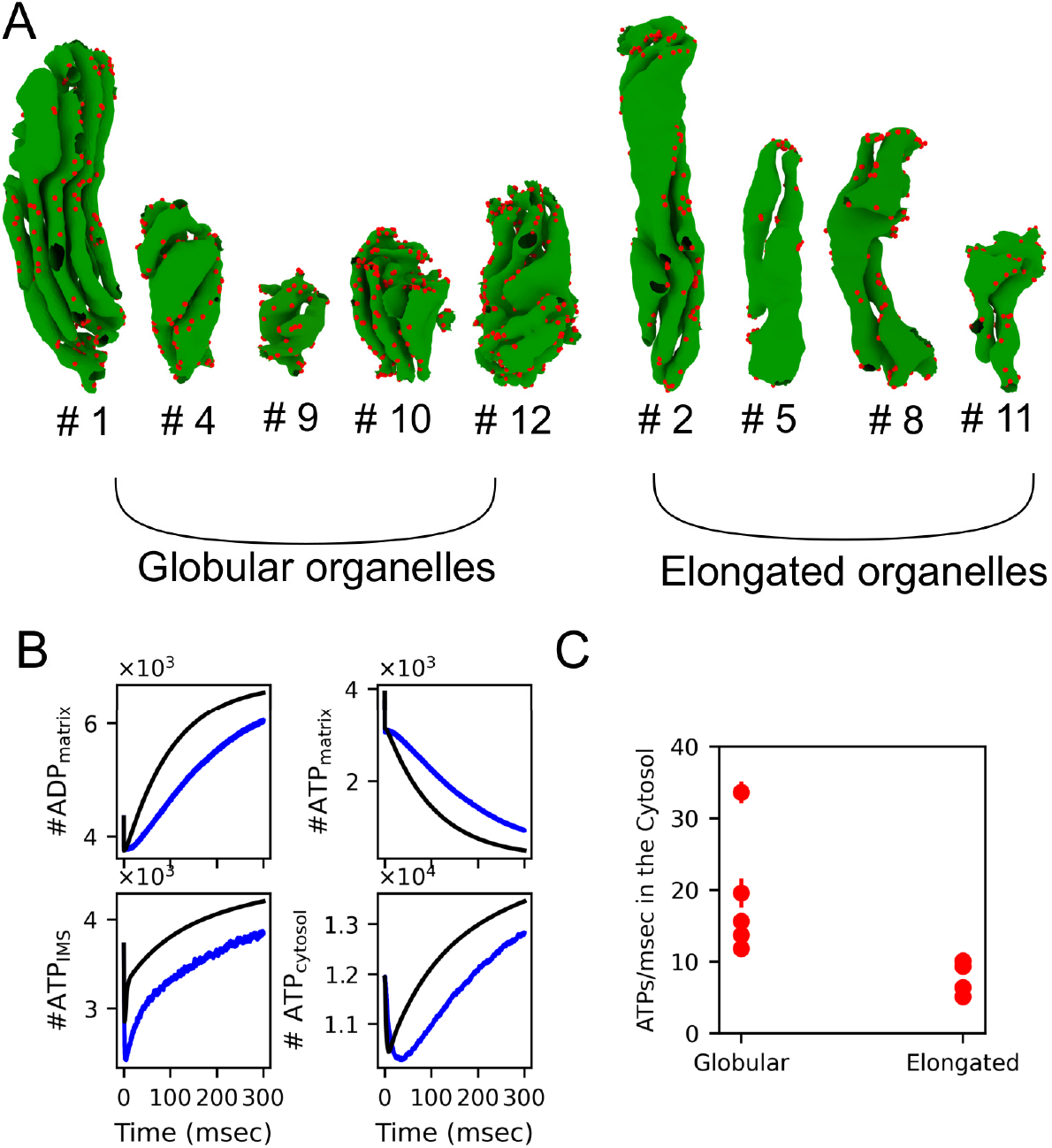
Spatial simulations: linking structure to function. A) Cristae membrane of the nine mitochondrial reconstructions considered with ATP synthases (in red) distributed in the areas of high curvature. B) Comparison of the ODEs simulations with the average traces of the spatial simulations for elongated mitochondrial reconstruction #2. C) ATP production rate in the cytosol for all reconstructions grouped accordingly to the structural features.

Using tools from differential geometry, we recently analyzed structural features of these reconstructions and based on this analysis we group them in two categories: globular organelles and elongated ones (Mendelsohn et al., 2021) (Figure 8A). Organelles in the globular group have larger surface area formed by vertices with high curvature. Large curved surface areas correspond to more number of ATP synthases in these reconstructions.

We performed spatial simulations for all reconstructions and we measured the ATP production rate in the cytosol for each reconstruction. For that, we performed a linear regression to single traces of the number of ATP molecules in the cytosol over time. Averaged rates are shown in Figure 8C. As expected from our analysis the ATP production rate in the cytosol is larger for globular organelles, which have larger areas of high curvature.

## 5. Discussion

Given the indispensable nature of mitochondria in cellular energy production, having accurate models of mitochondrial dynamics is critical for understanding their impact on cellular function.

In this work, we focused on developing a thermodynamically consistent model of ATP production in mitochondria to ensure the model is physically plausible. In doing so, we derived the relationships for the dependence of kinetic parameters on the membrane potential, proton and substrate concentration.

We explored different parameter regimes to understand in which conditions ATP production or its export are the limiting steps in making ATP available in the cytosol. Under the conditions considered, we found two regimes: at a low ratio of ANTs to ATP synthases, the export of ATP is limited by the ANTs. At ratios higher then 20:1, the ATP production becomes the limiting step. Similarly, the number of VDACs does not limit the export of ATP, unless there are very few VDACs present (in the order of 100s, for the ratio of ANTs/ATP synthases considered).

Finally, we performed spatial simulations on nine mitochondrial reconstructions derived from electron microscopy data. According to our parameter exploration, the number of ANTs and VDACs in these nine reconstructions are such that the synthesis of ATP is the limiting step on the ATP production rate in the cytosol and not its export. We further measured the rate for the same conditions, and relate this quantity to structural features of the organelles. We had mapped features of the OM to features of the IM recently in Mendelsohn et al. 2021 with tools from differential geometry and showed that globular organelles —characterized by smaller aspect ratio and smaller average first principal curvature of the OM— have larger curved CM surface areas. These organelles also have other features of high energy capacity like large crista shape factor and a high number of crista junctions per OM surface area. Others have shown that mitochondria at high-energy demanding locations in outer hair cells of young mice (Perkins et al., 2020) as well as mitochondria from high-spiking interneurons (Cserép et al., 2018) show a large crista shape factor and a high number of crista junctions per OM surface area. These features are also characteristic of our globular set, which suggests a high level of metabolic activity for such mitochondria. Here, using our mathematical model we found globular organelles generate more ATP in the cytosol than the elongated morphologies (Figure 8C).

The maximum rate measured with our spatial model is roughly 30 ATPs/ms (Figure 8C), this corresponds to 170 µM s^−1^ (considering the volume of the cytosol). Recent measurements in myocytes (Wescott et al., 2019) estimated the ATP production rate in the range 200-600 µM s^−1^ depending on the calcium concentration. The organelles considered in this manuscript are axonal and presynaptic reconstructions and are small (in the range of 0.037 to 0.012 µm^3^) compared to organelles in myocytes (in the order of 0.5 µm^3^ (Laguens, 1971)). This could explain our smaller ATP production rate in the cytosol.

We acknowledge that our modeling approach has a number of limitations. First, the membrane potential in our simulations is constant, we emulate a constant state of ATP production. Second, the ATP synthase flux considered in our model saturates sigmoidally at a large membrane potential (Figure 3B), as it has been shown for P. modestum bacteria and E. coli (Kaim and Dimroth, 1999), and is more sensitive to changes in the membrane potential than to ΔpH. However, recent experiments on heart and skeletal mitochondria (Wescott et al., 2019) show no saturation of the ATP synthase flux when increasing the membrane potential. Our model does not reproduce this behavior. Its voltage-dependence is a consequence of the detailed dynamic of ATP synthase model (Pietrobon and Caplan, 1985), which is drawn from physical assumptions about the rate constants, the membrane potential, and the pH concentration. Taking different assumptions or expanding the number of states could perhaps give a different flux membrane potential dependence. Third, the Mg^2+^-bound forms of ADP and ATP are the main substrate of ATP synthase in mitochondria. Since our goal was to generate a spatial model of a mitochondrion and Mg^2+^ is in large concentration in the matrix, we considered the free form of ADP as the main substrate for ATP production. This is an approximation, due to the limitations of using an agent-based model. However, this approximation has been taken in many mitochondrial models in the past (Magnus and Keizer, 1997; Bertram et al., 2006; Cortassa et al., 2003) and it is accepted in the community (Gnaiger, 2020).

Different approaches can be taken to model the spatial organization of mitochondria. From continuum approximations (Leung et al., 2021; Afzal et al., 2021) to discrete descriptions as we considered in our spatial simulations (with an agent-based reaction-diffusion simulator). Well-mixed continuum approximations based on concentrations like those in (Leung et al., 2021; Afzal et al., 2021) break down at low particle numbers. A discrete particle-based approach, as we considered in our spatial simulations with an agent-based reaction-diffusion simulator, is more accurate in this scenario (Bartol et al., 2015). Continuum approaches could be more challenging to integrate in complex geometries. Since we want to explore the impact of the complex mitochondrial morphology on ATP production and include the specialized location of ATP synthases at the rim of the cristae we opted for a discrete description in our spatial simulations.

These different approaches can lead to similar results. For instance, in the past we observed gradients of ATP with our discrete approach (Garcia et al., 2019) and so did (Afzal et al., 2021) with a continuum description. Our earlier version of the model (Garcia et al., 2019) is based on the same published data (Pietrobon and Caplan, 1985; Metelkin et al., 2006; Magnus and Keizer, 1997). The major improvement of the current version is that the parameters were constrained by thermodynamic considerations (Hill, 1989). The results, however, are consistent (in Supplementary Figure 5 we compare for reconstruction # 1 both versions of the model).

Finally, our results have implications for our understanding of disease states. For instance, near total loss of ATP synthase dimerization caused by a neurodegenerative mutation generated profound disturbances of mitochondrial crista ultrastructure in neurons derived from a Leigh syndrome patient (Siegmund et al., 2018). The main architectural feature different than in control cells is an increment in the angle of curvature at the tips of the cristae. This feature relates to our curvature measurement of the CM, which directly relates to the total number of ATP synthases. Altered mitochondrial structures have also been shown in aged animals (Perkins et al., 2020; Glavis-Bloom et al., 2023). Further investigation using the methods developed here will allow a deeper understanding of potentially impaired mitochondrial mechanisms in disease states.

## 6. Data Availability

All the material associated with this work is available in the repository: https://github.com/RangamaniLabUCSD/spatial_mito_model

## 7. Acknowledgments

This work was supported by Air Force Office of Scientific Research (AFOSR) Multidisciplinary University Research Initiative (MURI) grant FA9550-18-1-0051 (to P. Rangamani) and Office of Naval Research N00014-20-1-2469 (to P. Rangamani). All authors declare no known or potential conflict of interest including any financial, personal, or other relationships with other people or organizations that could inappropriately influence, or be perceived to influence their work.

## Abbreviations

ATP: adenosine triphosphate
ADP: adenosine diphosphate
OM: outer membrane
IM: inner membrane
ETC: electron-transport chain
Pi: inorganic phosphate
CM: cristae membrane
IMS: intermembrane space
VDAC: voltage dependent anion channel
ANT: adenine nucleotide translocator
ODEs: ordinary differential equations

## Footnotes

1 These calculations were reported to be in a Ph.D. thesis we could not get access (Magnus, 1995). Recently, we found part of these calculations were included in another PhD thesis (Balbir, 2007)

