## Supplementary Material for "Mitochondria morphology governs ATP production rate"

### 1 ATP/ADP translocator: Experimental results

The ATP/ADP translocator model is based on the work of Metelkin et al. 2006, and two additional states were added to track futile translocation in the spatial model. Starting from steady-state flux parameters from Metelkin et al. 2006 for ANTs from heart mitochondria (Kraemer and Klingenberg, 1982), we first estimated the parameters for the spatial implementation and the corresponding ODE model. To adapt the model to the non-steady state condition, the backward and forward rate constants were estimated from the dissociation constants. The forward rate constants were set as smaller than the diffusion-limited rate and the backward rate constants were set to satisfy the dissociation constant ratio. For instance, the dissociation constants at a membrane potential of 180 mV is $K_{Ti} \approx 400\,\mu M$ in the original paper, and in our simulations, 500 µM was chosen. Similarly, $K_{Di}$ was modified from 51 µM in the original paper to 25 µM in our simulations. We also estimated the dissociation constants from the matrix side $K_{Tm}$ and $K_{Dm}$ as 6.25 mM and 10 mM, respectively (the ratio is $\approx 1$). With these modifications, we were able to qualitatively reproduce the independent data from (Kraemer and Klingenberg, 1982; Duyckaerts et al., 1980) (Figure 1 and 2).

We also qualitatively reproduced the experimental data of Duyckaerts et al. 1980. Duyckaerts et al. 1980 proposed a mechanism for the ANT translocator which implies the formation of a ternary complex and proposed the explicit form for the flux as

$$v_o^{-1} = \frac{K_{AB}}{V}\frac{1}{[A][B]} + \frac{1}{V}. \tag{1}$$

From this relation, it follows that the reciprocal of the initial rate of exchange as a function of the concentrations of ADP in the matrix and outside should be linear as shown by the straight line in Figure 2E. From this relation, they also estimated the parameters from their data and obtained a value for $K_{AB}$ of $26.5\times10^{-3}\,mM^2$. This value is in good agreement with our simulations, which give a parameter value $K_{AB}$ of $27.06\times10^{-3}\,mM^2$ (Figure 2F).

### 2 ATP/ADP translocator: comparison with other models

We compare the ANT model with another model in the literature. With the ANT model implemented by Magnus and Keizer 1997, developed by Bohnensack 1982 we analyzed how the ANT flux depends on the $ATP_m/ADP_m$ ratio, and on the $ATP_i/ADP_i$ (Figure 3). Similar dependencies were shown in the main text for the ANT model implemented in this paper (Figure 4A-B in the main manuscript).

### 3 Spatial Simulations

#### 3.1 Same number of ATP synthases in all the reconstructions

We performed spatial simulation with the same conditions considered in the main text, but keeping the number of ATP synthases constant in all reconstructions (Figure 4). The first five reconstructions are in the globular set and the last four are in the elongated one. On average the globular set generates a rate of 17 ATPs/ms and the elongated one of 10 ATPs/ms, the difference is statistically significant (p-value $< 1.8\times10^{-6}$).

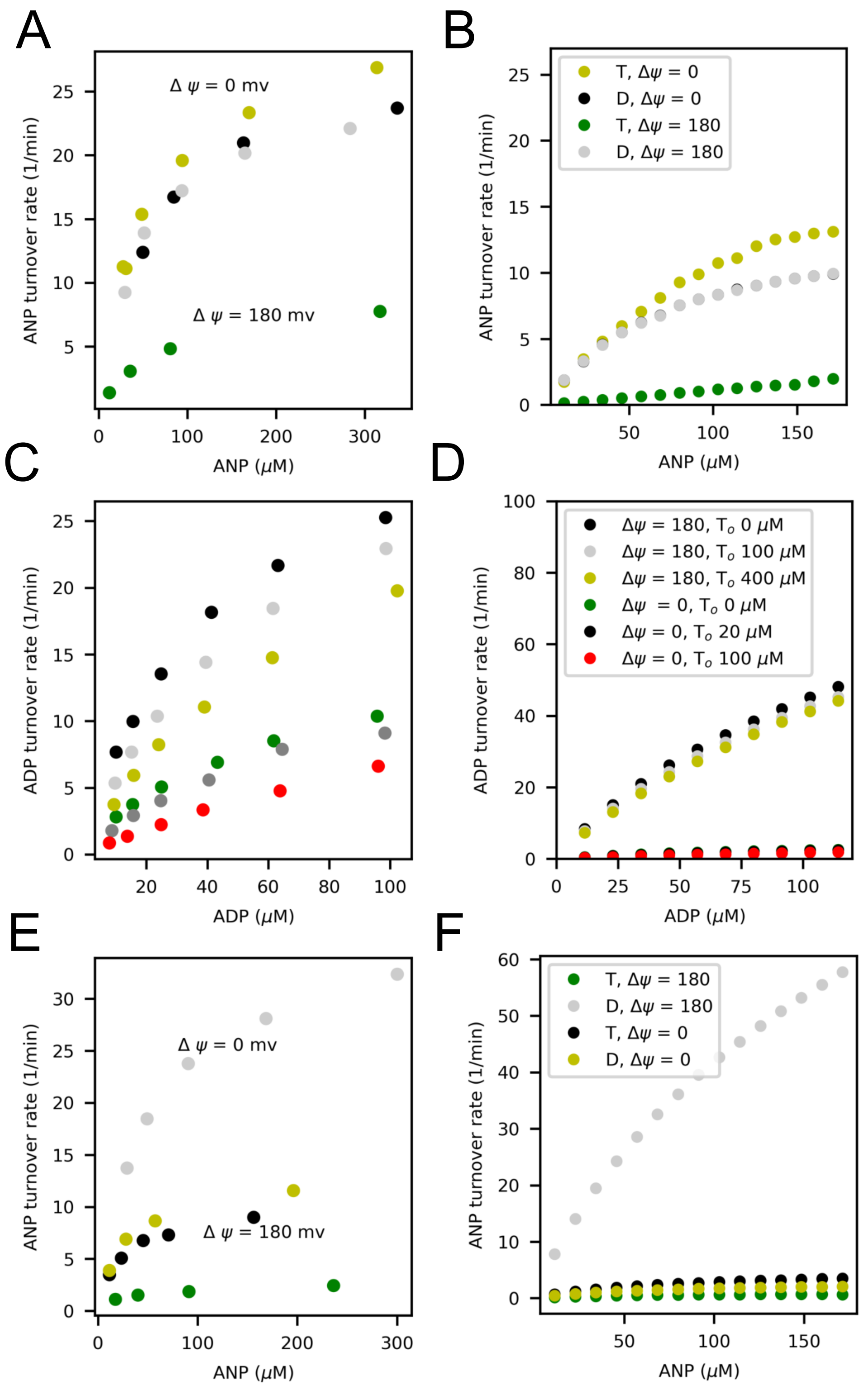


Figure 1: Qualitative reproduction of experimental fluxes from published work (Kraemer and Klingenberg, 1982). (A-B) ATP uptake rate vs external ATP concentration for a membrane potential ($\Delta\Psi$) of 0 and 180 mV in yellow and green, respectively. Black and grey points describe the ADP uptake rate vs external ADP concentration for a membrane potential of 0 and 180 mV. A) Transformed data from Kraemer et al.,(1982) (we followed the same approach as in Meletkin et al.) and (B) Our simulations, the flux was computed calculating an average over 30 seconds. ANP stands for ATP for the yellow and green points, and for ADP for Black and grey points. (C-D) ADP uptake rate vs external ADP concentration for membrane potential ($\Delta\Psi$) of 180 mV and ATP concentrations outside the liposomes ($T_o$) of 0, 100, 400 µM shown in black, light grey and yellow points, respectively. Green, grey, and red points describe the dependency with the membrane potential of 0 mV and concentrations of ATP concentrations outside the liposomes ($T_o$) of 0, 20, 100 µM, respectively. The concentrations of ADP and ATP inside the liposomes were 5 m. (C) ATP uptake rate vs ATP concentration, yellow and green points for a membrane potential ($\Delta\Psi$) of 0 and 180 mV respectively, and ADP concentration equal to the ATP concentration. ADP uptake rate vs ADP concentration, black and grey points for a membrane potential of 0 and 180 mV respectively, and ATP concentration equal to the ADP concentration.

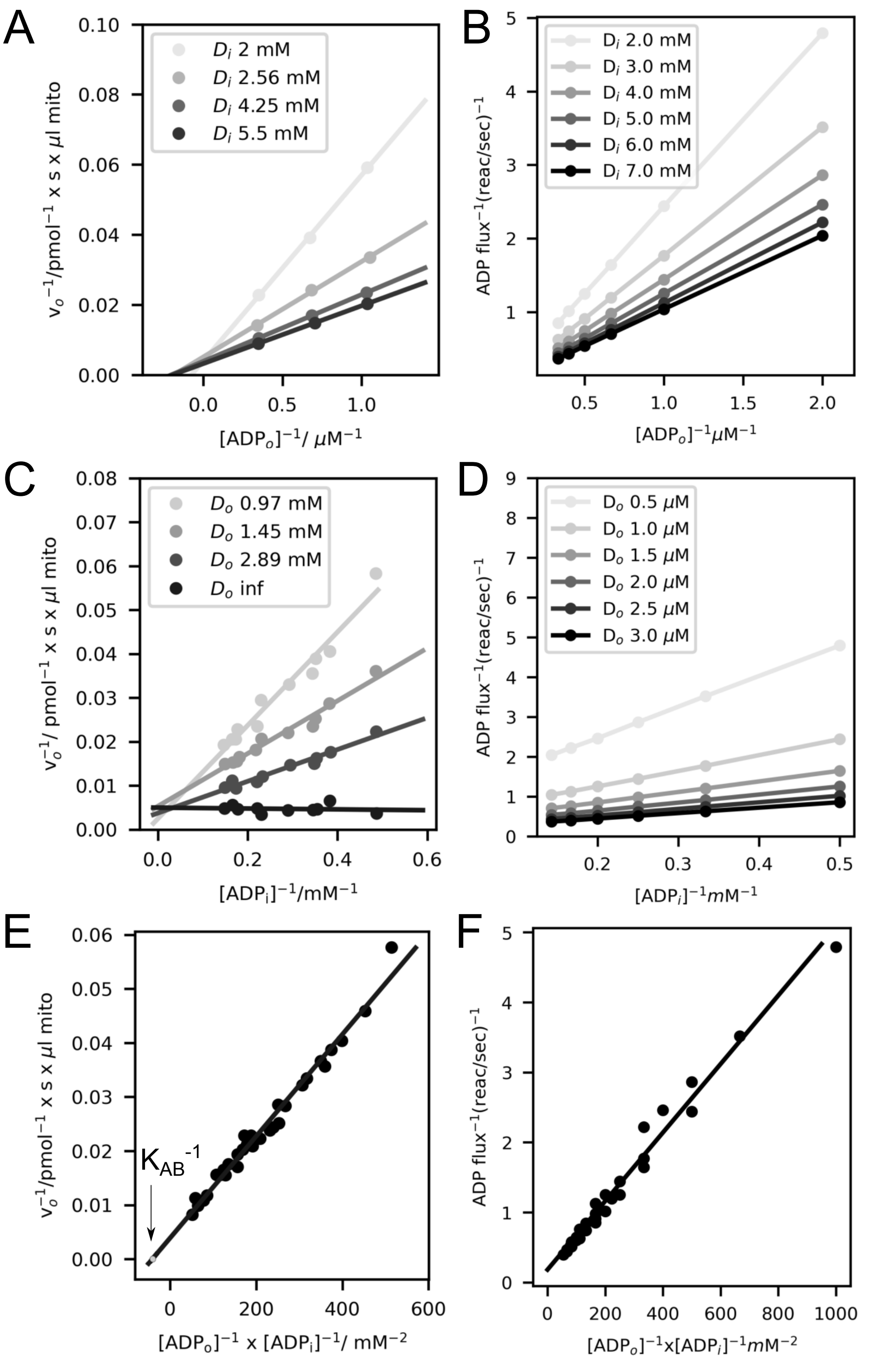


Figure 2: Qualitative reproduction of the experimental results from published work (Duyckaerts et al., 1980). (A-B) Reciprocal of the initial rate of ADP exchange as a function of the reciprocal of the external ADP concentration. The initial rate was measured for three external-ADP concentrations and eleven internal-ADP concentrations (only four are shown). The lines were calculated by the method of least squares. (A) Experimental data reproduced from Duyckaerts et al. 1980 and (B) our simulations. (C-D) Reciprocal of the initial rate of ADP exchange as a function of the reciprocal of the internal-ADP concentration. The initial rate was measured for three external ADP concentrations and eleven internal-ADP concentrations. (C) Data reproduced from Duyckaerts et al. 1980 and (D) our simulations. (E-F) Reciprocal of the initial rate of ADP exchange as a function of the reciprocal of $[ADP]_i[ADP]_o$. (E) Data reproduced from Duyckaerts et al. 1980, the parameter value $K_{AB}$ is $26.5 \times 10^{-3}\,mM^2$ and (F) our simulations, our estimation of $K_{AB}$ from our simulations is $27.06 \times 10^{-3}\,mM^2$.

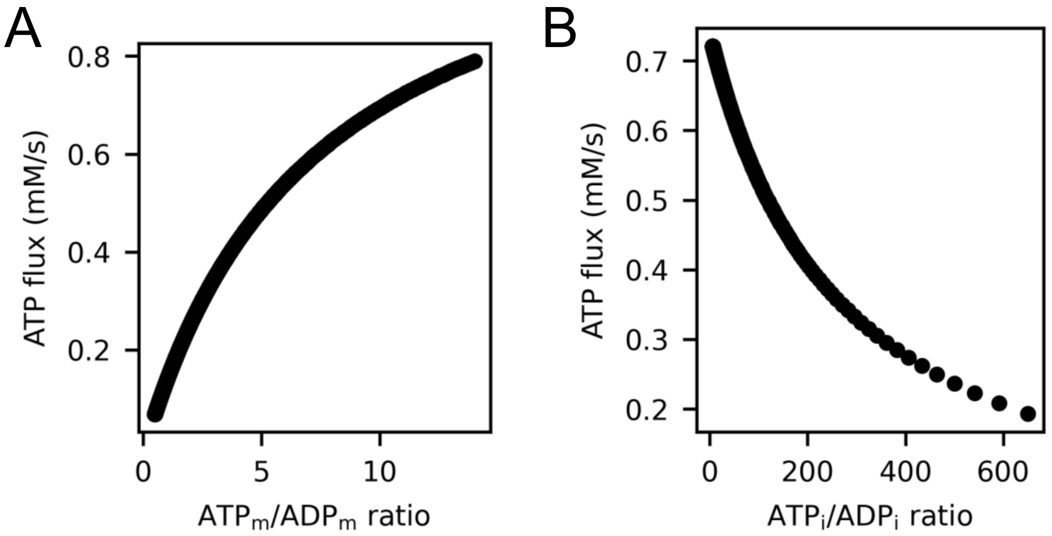


Figure 3: Comparison with ANT model developed by Bohnensack 1982. (A) ATP flux dependence on the ATP to ADP ratio on the matrix. (B) ATP flux dependence on the ATP to ADP ratio in the IMS at a constant membrane potential of 180 mV.

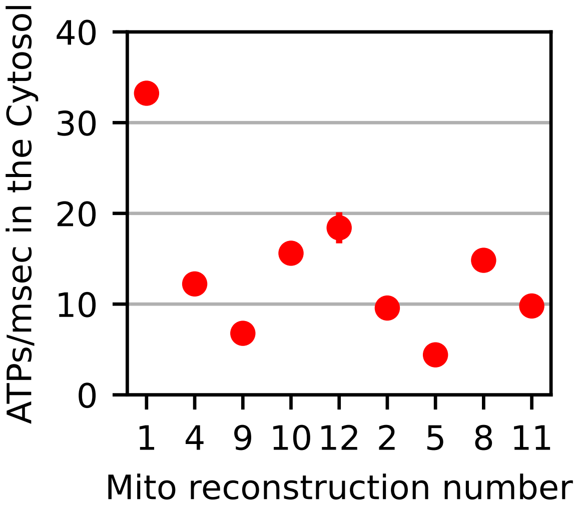


Figure 4: ATP production rate in the cytosol for all reconstructions, considering the same number of ATP synthases (204). The first five reconstructions belong to the globular set and the last four to the elongated one. On average the ATP production rate is also higher for the globular set under these conditions.

#### 3.2 Comparison to our previous version of the model

We compare the dynamics of our previous version of the model (Garcia et al., 2019) and the current version for the mitochondrial reconstruction number 1 (Figure 5). The ATP production rate with the new parameters of the model is 34 $\pm$ 2 ATPs/ms whereas, with the previous version, the rate was 68 $\pm$ 2 ATPs/ms for the same reconstruction.

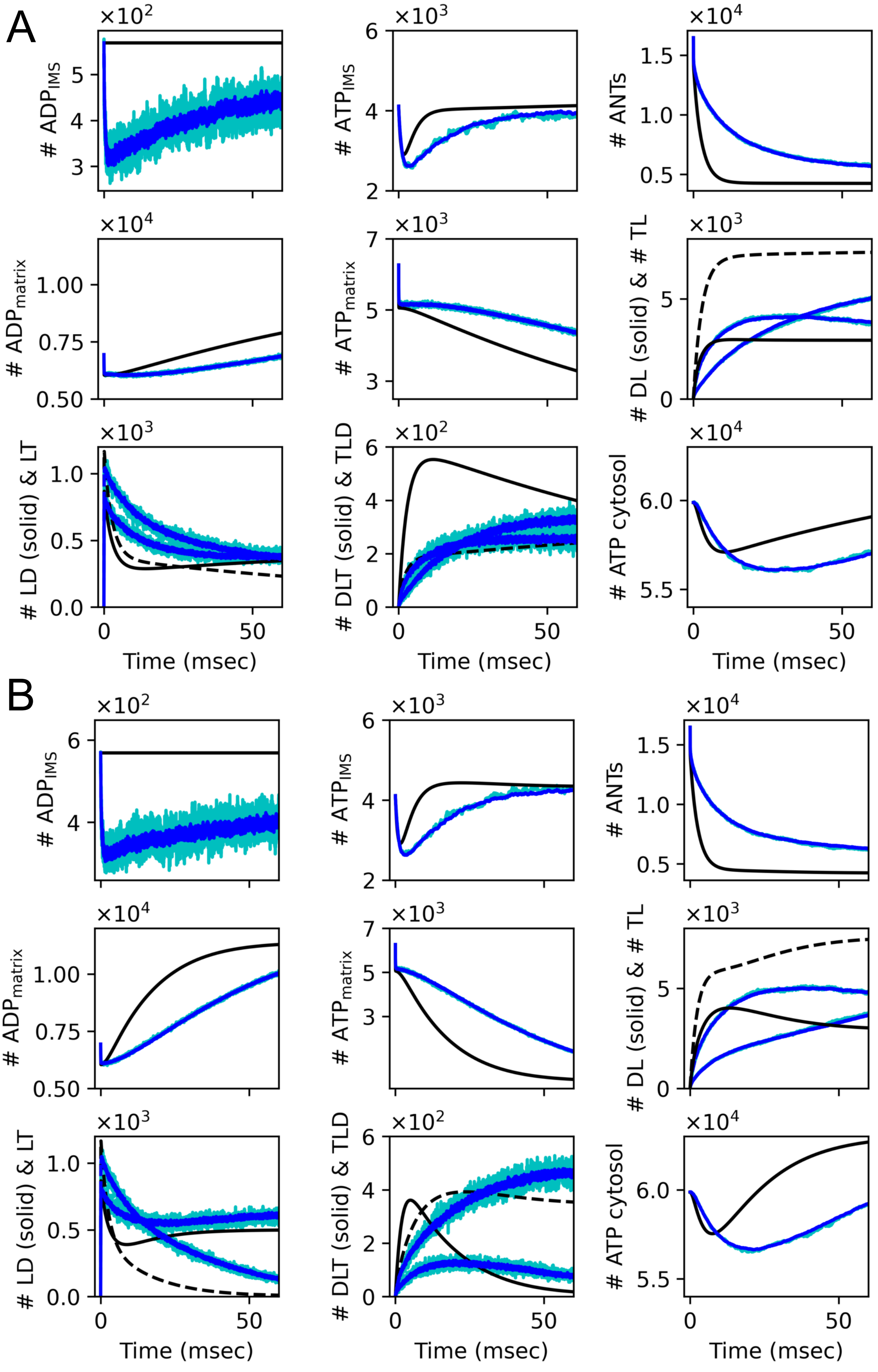


Figure 5: Comparison to our already published model (Garcia et al., 2019). Average traces of the number of molecules in the different compartments (in blue), a single trace (in cyan) and results of the ODEs (in black) on the left (panel A) results of our current version of the model and on the right (panel B) results for the same conditions for the old version of the model (Garcia et al., 2019), for reconstruction #1.
